# Predicting subclinical psychotic-like experiences on a continuum using machine learning

**DOI:** 10.1101/380162

**Authors:** Jeremy A Taylor, Kit Melissa Larsen, Ilvana Dzafic, Marta I Garrido

## Abstract

Previous studies applying machine learning methods to psychosis have primarily been concerned with the binary classification of chronic schizophrenia patients and healthy controls. The aim of this study was to use electroencephalographic (EEG) data and pattern recognition to predict subclinical psychotic-like experiences on a continuum between these two extremes in otherwise healthy people. We applied two different approaches to an auditory oddball regularity learning task obtained from *N* = 73 participants:

1. A feature extraction and selection routine incorporating behavioural measures, event-related potential components and effective connectivity parameters;
2. Regularisation of spatiotemporal maps of event-related potentials.

Using the latter approach, optimal performance was achieved using the response to frequent, predictable sounds. Features within the P50 and P200 time windows had the greatest contribution toward lower Prodromal Questionnaire (PQ) scores and the N100 time window contributed most to higher PQ scores. As a proof-of-concept, these findings demonstrate that EEG data alone are predictive of individual psychotic-like experiences in healthy people. Our findings are in keeping with the mounting evidence for altered sensory responses in schizophrenia, as well as the notion that psychosis may exist on a continuum expanding into the non-clinical population.

## 1. Introduction

It has been theorised that psychotic experiences are distributed along a continuum. On the lower end of this scale, schizotypy, or ‘psychotic-like’ experiences, can also occur within the healthy population (Verdoux and van Os, 2002). These psychotic-like experiences are considered a potential risk factor for schizophrenia, particularly as those experiencing such phenomena are at higher risk of developing schizophrenia spectrum disorders (Wang et al., 2018). Whilst delineating the brain mechanisms underlying the continuum of psychosis in the healthy population could potentially help in understanding the disease trajectory, individuals with subclinical psychotic-like experiences are not ‘help-seeking’ and therefore have not been clinically assessed against the criteria of ‘clinical high-risk’ or assessed for meaningfully attenuated positive symptoms (Cicero et al., 2014).

Multivariate machine learning regression (Cohen et al., 2011; Janssen et al., 2018; Madsen et al., 2018) is a pertinent means for modelling psychosis on a continuum and making predictions at the individual level. Machine learning algorithms can automatically detect patterns within a given dataset and train computational models to make such predictions about new test data, based on prior observations from the training data. Whilst these predictions can be either categorical or continuous in nature, previous machine learning studies in the field of psychiatry have primarily been concerned with the binary classification of patient groups and healthy controls. Stephan et al. (2017) argued against this dichotomy and questioned the utility of models capable of reproducing discrete, predefined categories. In recent years, there has been a shift away from these formal diagnostic criteria toward a dimensional approach as per the Research Domain Criteria (RDoC; Cuthbert and Insel, 2013). In order to model the complete spectrum of possible disease states which span from normal through to abnormal, it is equally important to asses those between these two extremes, such as psychotic-like experiences in otherwise healthy individuals.

Whilst machine learning methods have previously been used to predict continuous measures of personality traits such as anxiety (Portugal et al., 2019), negative affect (Fernandes Jr et al., 2017) and emotional dysregulation (Portugal et al., 2016), thus far it remains unclear whether the severity of psychotic-like experiences can be predicted from neuroimaging data when treated as a continuous variable (Madsen et al., 2018). There have however been several recent studies focusing on binary classification. For example, Jeong et al. (2017) categorised individuals into high and low schizotypy, classifying between these subgroups based on event-related potentials from an emotional perception paradigm. Kalmady et al. (2020) applied an existing schizophrenia *vs*. control classifier to resting state functional magnetic resonance imaging (fMRI) data from unaffected first-degree relatives and found that false positives (i.e. misclassified as schizophrenia patients) had higher schizotypal personality traits than true negatives. Similarly, Krohne et al. (2019) classified between high and low measures of social anhedonia within a schizotypy group using a theory of mind and empathy social cognition task in fMRI. Zarogianni et al. (2017) have also used schizotypy scores in combination with other neuroanatomical features in predicting schizophrenia onset. Finally, post hoc analyses by Di Carlo et al. (2019) suggested that binary misclassifications of healthy controls as schizophrenia patients from structural images of the thalamus were related to higher schizotypy.

The aim of this study was to predict a range of psychotic-like experiences on a continuum within a healthy population using machine learning regression methods on electrical brain activity, recorded non-invasively during an oddball task. Oddball tasks elicit a robust mismatch negativity response (MMN), which can be measured using electrencephalography (EEG) and is consistently shown to be reduced in schizophrenia (Todd et al., 2012), and predictive of people at high risk transitioning into schizophrenia (Bodatsch et al., 2011). Given the significant translational potential of MMN in schizophrenia, we hypothesized that it would be predictive of subclinical psychotic-like experiences in a community sample. To test this, we applied two approaches:

1. A pipeline incorporating sequential feature selection from a range of behavioural measures, event-related potential components, as well as effective connectivity between brain regions, providing us with a high level of model interpretability and enabling us to better determine the importance of individual features across multiple subdomains;
2. Applying a spatiotemporal approach to event-related potentials as to determine which particular responses are most predictive of psychotic-like experiences, and within those responses, which spatial locations and time points contribute most to the overall predictions.

In so doing, we hoped to provide further evidence that subclinical psychosis may exist on a continuum within the healthy population.

## 2. Methods

### 2.1. Participants

The results presented in this paper were obtained by merging two datasets and repeating a previous analysis protocol initially performed on a smaller sample. For full transparency, the results from our initial sample of *N*_1_ = 31 are available in supplementary materials (Table S1 and Figures S2 to S4). An additional *N*_2_ = 45 were recruited, of which three were excluded due to instrumentation failure and poor signal. We have accounted for these models in our corrections for multiple comparisons.

The combined group comprised a total of *N* = 73 high functioning individuals with no prior history of psychiatric or neurological disorders (age 18 to 39 years, mean = 24.51 years, *SD* = 4.94, 36 males). A screening process preceding recruitment confirmed that all participants had no history of psychiatric or neurological disorders, were not currently taking medication, and had not used any illicit drugs in the past three months. All participants provided written informed consent and were paid for their participation. The study received approval by the University of Queensland Human Research Ethics committee.

To obtain individual schizotypy scores, all participants completed the Prodromal Questionnaire (PQ; Loewy et al., 2005), which comprises of 92 items assessing four subscales; positive (mean ± *SD* = 28.53 ± 28.09, range 0-143), negative (17.95 ± 15.01, 0-62), disorganised and general symptoms (30.15 ± 22.79, 1-105). Responses recorded the frequency of psychotic experiences (‘never’, ‘once or twice per month’, ‘once per week’, ‘few times per week’, ‘daily’) with the total PQ (76.63 ± 63.68, 2-308) computed as the sum of all subscales.

### 2.2. Experimental design

EEG was recorded whilst participants listened to an auditory duration oddball paradigm with reversal learning task. The stimuli comprised streams of predictable ‘standard’ tones occurring at 80% probability interspersed with infrequent ‘deviant’ tones at 20% probability. All stimuli were presented at 500ms intervals with pure frequency of 500Hz and smooth rise and fall transitions of 5ms. The duration assigned to standard and deviant stimuli alternated between ‘short’ (50ms) and ‘long’ (100ms) tones at varying regularities within ‘stable’ and ‘volatile’ blocks as illustrated in Figure 1a. In a stable block, duration assignment is fixed throughout the block. In a volatile block, the duration which is more likely at first is occasionally ‘reversed’ to become less likely, with three probability reversals per block. Over the course of the experiment, a total 2000 stimuli were presented (1000 each for both short and long tones) over eight blocks (four stable and four volatile).

**Figure 1.**
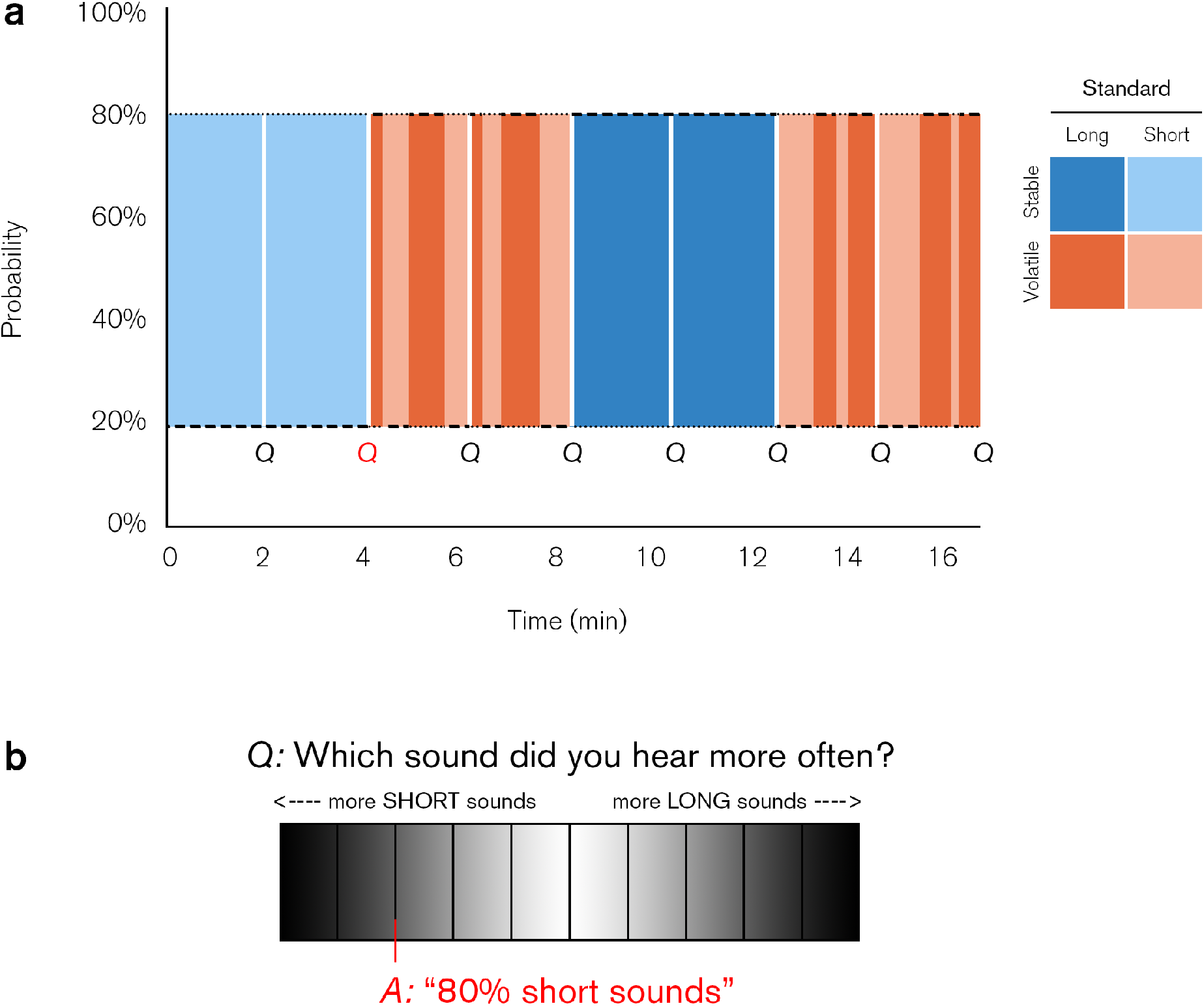
Schematic of reversal oddball task. **(a)** Exemplar stable (blue) and volatile (red) blocks comprising short (50ms, dots) and long sounds (100ms, dashes). Participants were asked to estimate the probability of sounds (denoted *Q*) every 2:08 minutes and rate their confidence. **(b)** Scale used for response input.

Participants were instructed to attend to these sounds and report the probability of the more frequent tones heard and level of confidence in their estimate at the end of each block via keyboard and mouse. Probability was reported on a linear scale (see Figure 1b), with the leftmost end of the scale corresponding to 100% short sounds and the right end 100% long sounds. Confidence was reported on an ordinal three-point scale (1 = ‘not confident’, 2 = ‘moderately confident’, 3 = ‘very confident’). For more detail regarding the task and stimulus paradigm, refer to Dzafic et al. (2020).

### 2.3. Data collection and pre-processing

#### 2.3.1. EEG

Continuous EEG data were recorded using a 64 channel Biosemi ActiveTwo system (Amsterdam, Netherlands) with electrode placement according to the international 10-20 standard (Oostenveld and Praamstra, 2001) at a sampling rate of 1024Hz. Offline signal processing was performed using SPM12 (www.fil.ion.ucl.ac.uk/spm) (Litvak et al., 2011). Data were referenced to the scalp average, downsampled to 200Hz, and bandpass filtered with a 0.5Hz high-pass Butterworth filter. Eye blinks were detected using the vertical electrooculogram channel and corrected using the Berg method (Berg and Scherg, 1994). Events were epoched using a −100 to +400ms peri-stimulus time window. Trials were thresholded for artefacts at 100µV, robustly averaged (Wager et al., 2005), low-pass filtered at 40 Hz, and baseline corrected between −100 and 0ms.

#### 2.3.2. Feature selection approach

To facilitate feature selection, a subset of 20 variables was extracted from the dataset; event-related potential (ERP) components, behavioural measures, and effective connectivity between brain regions. The definition of these features was motivated by previous literature (Randeniya et al., 2018).

A total of six ERP components for the standard, deviant and Mismatch Negativity (MMN; deviant − standard) responses under stable and volatile conditions were obtained from the frontocentral channel (Fz), identifying peak amplitude within the 150-250ms time window, i.e. the typical scalp location and latency associated with the prediction error responses observed in prior studies using oddball paradigms (Garrido et al., 2009).

Effective brain connectivity associated with the effect of environmental stability was estimated using Dynamic Causal Modelling (DCM) of a well-defined functional brain architecture (Garrido et al., 2007; Opitz et al., 2002), incorporating a fully-connected, three-level hierarchical brain model which underlies the generation of prediction error in auditory oddball paradigms. This model included: bilateral primary auditory cortices (A1; MNI coordinates: left [−42, −22, 7] and right [46, −14, 8]; the cortical input sources), bilateral superior temporal gyri (STG; left [−61, −32, 8] and right [59, −25, 8]), and bilateral inferior frontal gyri (IFG; left [−46, 20, 8] and right [46, 20, 8]) with individual structures modelled as separate dipoles for the two hemispheres. Nine model architectures were defined with varied combinations of forward and backward connections between the A1 and STG, IFG and STG, plus an intrinsic A1 connection resulting in a total ten possible connections. Bayesian model averaging was performed to estimate the strength of effective connectivity for each of these ten connections (referred to here as the DCM parameters, obtained from the *B* matrix) across all models (resulting from a weighted average of each connectivity parameter by the likelihood of the model under which that parameter was estimated). Note that the parameter estimation process was done separately for each participant without model selection, such that each training set remains completely independent from the testing dataset. A similar approach, referred to as generative embedding has been used previously (Brodersen et al., 2011), however in that paper, model optimization was done using the pooled data (both training and testing). For more detail regarding DCM fitting and parameter estimation, refer to Dzafic et al. (2020).

#### 2.3.3. Spatiotemporal approach

To facilitate spatiotemporal analyses, the ERPs for each participant, response and condition were converted to NIfTI images. At each time point, data were spatially interpolated between channels to form a two-dimensional scalp map, represented by a 32 × 32 matrix. These spatial images were then concatenated in temporal order, resulting in a three-dimensional spatiotemporal volume. Images were smoothed using a Gaussian filter with full width at half maximum of 12mm × 12mm × 20ms. In contrast with the feature selection approach, no further assumptions were made about channel locations or time windows.

### 2.4. Modelling

In all approaches described below, the target variables were the individual PQ scores, weighted by the frequency of occurrence and log-transformed to fulfil normality assumptions.

### 2.4.1. Feature selection approach

Regression modelling was performed on three sets of features extracted from behavioural, ERP and DCM data using the Huber regressor (Holland and Welsch, 1977; Huber, 1981) with an additional sequential feature selection process (Guyon and Elisseeff, 2003), reducing the dimensionality of the data from the complete set of *M* features, *X*_*M*_, to a subset of *m* features, where *m* ≤ *M*. Our pipeline, as illustrated in Figure 2, was implemented in the Scikit-learn library (version 0.22.1; Pedregosa et al., 2011) for Python (version 3.6.1; www.python.org) with sequential feature selection using the mlxtend library (version 0.17.1; rasbt.github.io/mlxtend).

**Figure 2.**
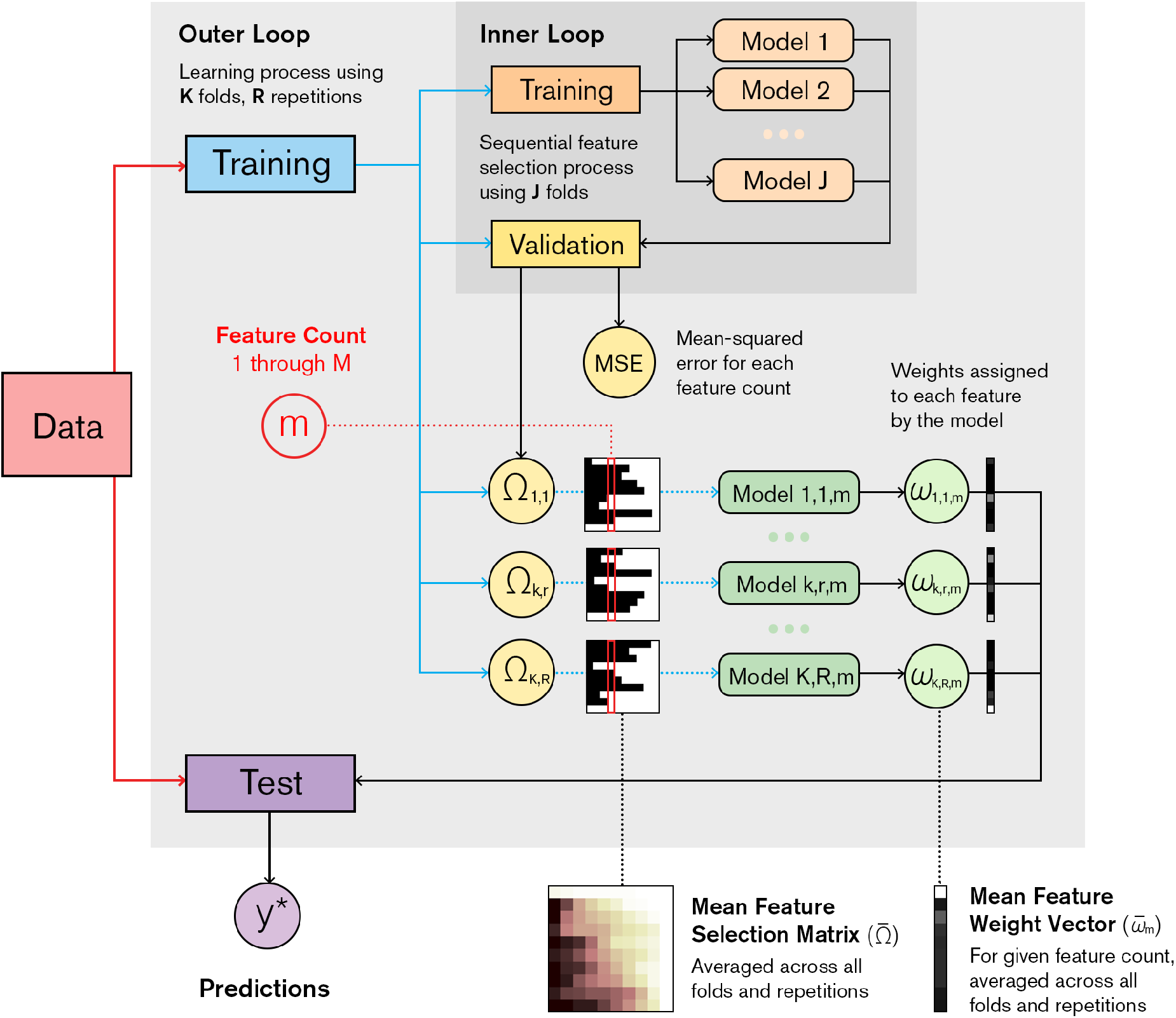
Schematic of machine learning pipeline. In the outer cross-validation loop, the collective dataset is partitioned into *K* folds, of which one is used for testing (purple) and the remaining set of *K*−1 folds for training (blue) a sub-model. This partitioning is repeated *R* times with differences in sampling. Feature selection is performed with an inner cross-validation loop, which partitions the training set from the outer loop into *J* folds, which form secondary training (orange) and validation sets (yellow). This process returns a binary matrix, Ω_*k,r*_, indicating the order in which features are selected. The optimal feature count, *m*, is that which achieves minimal mean-squared error across all folds and repetitions. There are a total *K* × *R* models (green), each of which are based on the *m*^th^ column of Ω_*k,r*_. The training process assigns weights, *ω*_*k,r,m*_, to each feature and are used to compute predictions, *y*\*, of independent samples in the test set. The mean feature selection matrix, 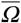, and mean feature weight vector, 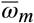, i.e. the average across all *K* folds and *R* repetitions, are used to intuit the feature importances.

Sequential forward selection was performed with a nested *J* = 5-fold cross-validation loop (shown in orange; Cawley and Talbot, 2010) on the training set from the main cross-validation loop (blue) to obtain the features which minimised mean-square error. We represent the order in which features are selected for a given fold, *k*, and repetition, *r*, as a binary *M* × *M* matrix, denoted here as *Ω*_*k,r*_, with individual features as rows, the feature count or algorithm iterations as columns, selected features as ones and unselected features as zeros. We enforce the criterion that all sub-models must share the same number of features and level of complexity, however, the differences in training data between folds and repetitions will result in different features being selected for each sub-model. The optimal number of features, denoted *m*, is that which achieved minimal mean-squared error on the validation set (yellow). From the *Ω*_*k,r*_ matrices, we extract a feature selection mask from the *m*^th^ column, determining which features form the basis for each sub-model. Through the training process, each of these features are assigned weights, denoted *ω*_*k,r,m*_. Each model then uses these weightings to make predictions, denoted *y*\*, of the test data (purple), which is strictly independent of both the optimisation and training processes. Using this pipeline, we specified four models; one based on each subset of the data, categorised as behavioural, ERP and DCM measures, as well as a combined model using all three measures. To summarise the frequency at which a given feature is selected, we average across all feature selection matrices 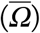, sorting features according to mean selection rate (i.e. the average across all columns) and in particular, inspect the optima from column *m*. To assess the weightings assigned to each feature upon selection, we also average across all weight vectors for the feature count 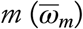.

### 2.4.2. Spatiotemporal feature sets

Regression modelling was performed using the kernel ridge regression algorithm (Hoerl and Kennard, 1970), as implemented within the Pattern Recognition for Neuroimaging Toolbox (PRoNTo, version 2.1; Schrouff et al., 2013). We used the default hyperparameters and the identical cross-validation scheme as in the feature selection approach. Ridge regression minimises the sum of squared residual errors whilst also penalising the size of coefficients through regularisation.

A total of four models were specified; one for each response (i.e. the standard, deviant and MMN difference waves), and a combined model concatenating all three images. For each model, we obtained mean feature weight maps, again averaging across all folds and repetitions. We compute an element-wise multiplication between the mean weight map and the grand mean activity associated with the model of interest, referred to henceforth as the ‘feature importance’ map (for illustration, refer to Supplementary Material, Figure S1).

#### 2.4.3. Evaluation of performance

Both mean square error (MSE), as well as the linear dependence between predictions and target variables, as measured using Pearson’s correlation coefficient (*R*), was used to evaluate model performance and identify any potential biases. The *p*-values for each model were computed through 1000 random permutations of the target PQ scores, with statistically significant results (*p* ≤ 0.05) for both measures suggesting that the algorithm had learnt some pattern within the data above chance (Golland and Fischl, 2003), subject to Bonferroni correction for multiple comparisons.

## 3. Results

The performance metrics for all models are summarised in Table 1. Overall, the spatiotemporal approach yielded greater performance in comparison with the feature selection procedure, of which the stable standard response demonstrated superior predictive performance (*R* = 0.42, MSE = 0.837, *p* = 0.032 corrected) in comparison with the deviant, MMN and combined feature sets (*R* = −0.04-0.44, MSE = 1.061-1.124). Equivalent models based on the volatile condition demonstrated markedly lower performance (*R <* 0, MSE = 1.123-2.029) and did not reach significance. The feature selection approach applied to the DCM connectivity parameters, ERP components and behavioural metrics did not identify a predictive pattern within these data and was non-significant.

**Table 1.** Summary of model performance. *p*-values are computed via 1000 permutations with Bonferroni correction for multiple comparisons. Significant results are highlighted in grey.

| Method | Features | R | R p-value<br>(uncorrected) | R p-value<br>(corrected) | MSE | MSE p-value<br>(uncorrected) | MSE p-value<br>(corrected) |
| --- | --- | --- | --- | --- | --- | --- | --- |
| Feature Extraction<br>and Selection | Behavioural | 0.167 | 0.050 | 0.400 | 1.000 | 0.049 | 0.392 |
|  | ERP Components | -0.314 | 0.696 | 1.000 | 1.085 | 0.676 | 1.000 |
|  | DCM Parameters | 0.113 | 0.141 | 1.000 | 1.041 | 0.124 | 0.992 |
|  | Combined | 0.170 | 0.089 | 0.712 | 1.035 | 0.081 | 0.648 |
| Spatiotemporal<br>Images | Stable Standard | 0.417 | 0.002 | 0.032 | 0.837 | 0.002 | 0.032 |
|  | Stable Deviant | 0.079 | 0.042 | 0.672 | 1.089 | 0.858 | 1.000 |
|  | Stable MMN | -0.039 | 0.220 | 1.000 | 1.124 | 0.988 | 1.000 |
|  | Stable Combined | 0.440 | 0.002 | 0.032 | 1.016 | 0.045 | 0.720 |
|  | Volatile Standard | -0.003 | 0.141 | 1.000 | 1.123 | 0.991 | 1.000 |
|  | Volatile Deviant | -0.143 | 0.516 | 1.000 | 1.188 | 1.000 | 1.000 |
|  | Volatile MMN | -0.199 | 0.695 | 1.000 | 1.179 | 1.000 | 1.000 |
|  | Volatile Combined | -0.018 | 0.448 | 1.000 | 2.029 | 1.000 | 1.000 |

Observed and predicted PQ scores for the stable standard spatiotemporal model are shown in the prediction plot in Figure 3. Although this model has a mean square error of 0.837 (equivalent to 17% error on average), there is a lower margin for error on the mid to high-range band of the spectrum, whereas we can observe that error is high on the lower end of the spectrum. This may suggest a possible bias towards the sample mean and undersampling of these ultra-low scores.

**Figure 3.**
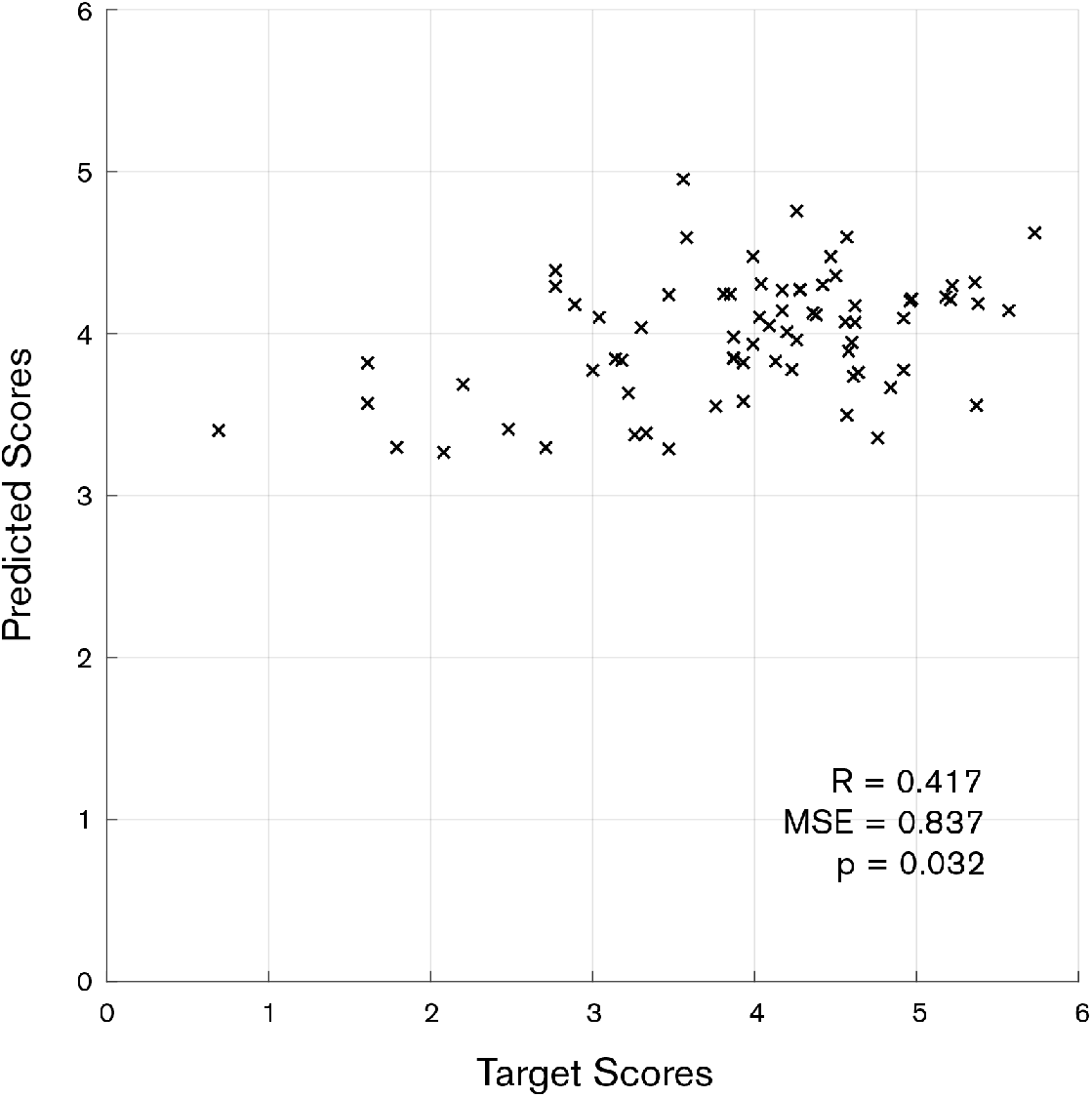
Plot of model predictions and true observed PQ scores, based on spatiotemporal stable standard ERP feature set, using kernel ridge regression and regularisation. All scores are normalised via log transform.*p*-values were computed via 1000 permutations with Bonferroni correction for multiple comparisons.

To intuit which features are driving the model predictions (shown in Figure 3), we computed the feature importances as shown in Figure 4, with positive importances contributing to a higher PQ and negative importance contributing to lower PQ (Taylor and Garrido, 2020). For the purposes of visualisation, we threshold the top 5% of both positive and negative importances in Figures 4A and 4B, respectively. Note, however, that although the majority of features have zero or low importance, all voxels in the image contribute towards the model predictions. The distribution of all signed feature importances within the volume is shown in Figure 4C. The main clusters of voxels contributing most to lower scores (Figure 4A) occur in central channels at approximately 50ms and between 150 and 275ms, whereas those contributing most to high scores (Figure 4B) occur between approximately 75 and 150ms over the frontal and left posterior channels.

**Figure 4.**
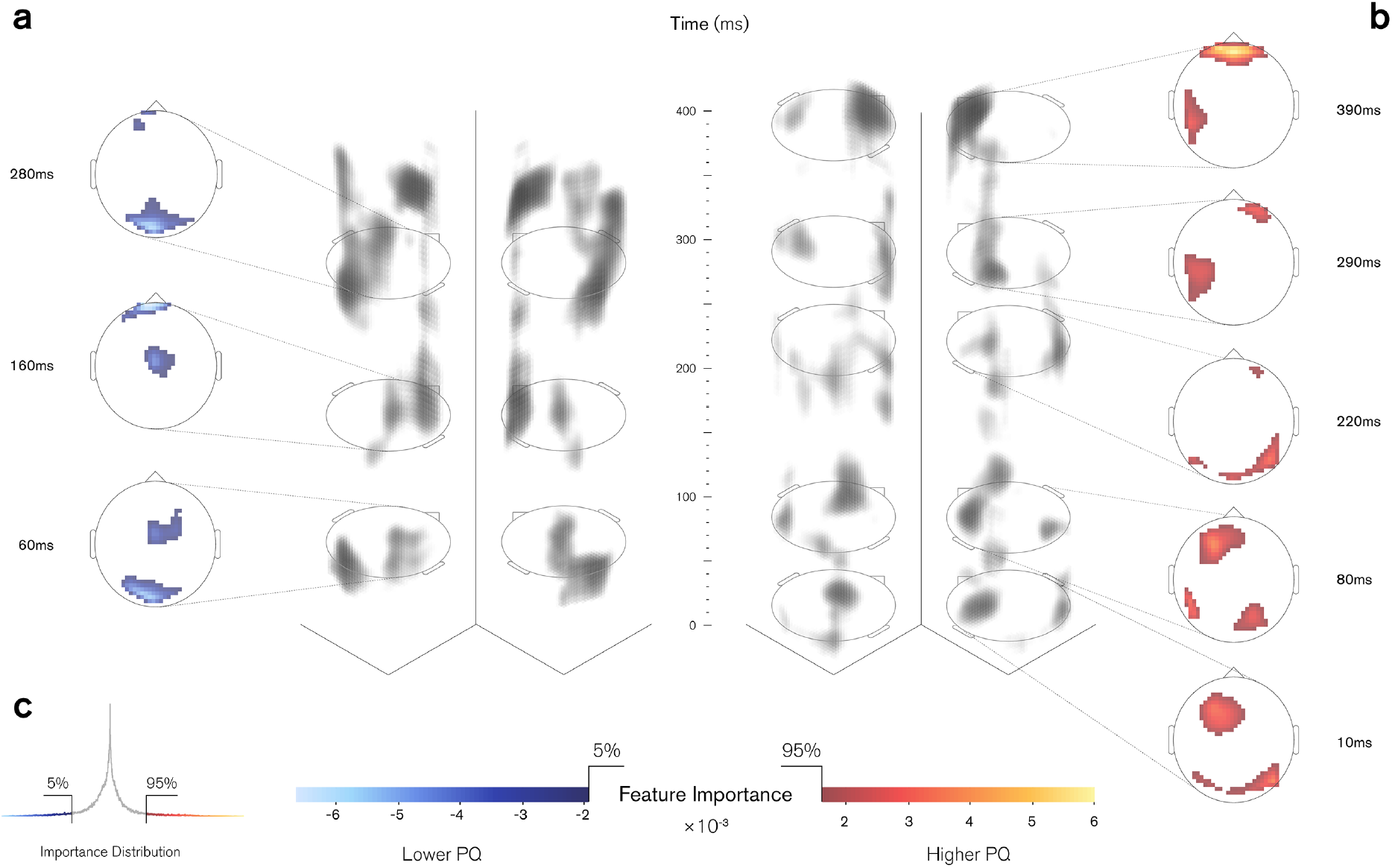
Spatiotemporal features of greatest contribution to predictions of PQ scores in stable standard response model. **(a-b)** Full feature set input to the model is rendered as a 3D volume (grey structures) where the spatial scalp dimensions are shown on *x-y* plane and, time domain along vertical *z-*axis. These grey structures represent the clusters of activity, or subgroups of features, with the greatest importance. Transparency of each individual voxel is mapped to the net contribution, with darker clusters indicating greater contribution. For the main clusters (cool and warm colormaps adjacent to grey structures), 2D scalp map annotations refer to time and location of peak activity for **(a)** low and **(b)** high PQ predictions, shown in cool and warm colormaps, respectively. Voxels are thresholded at top 5% of negative and positive contributions respectively. Each volume is displayed from two different angles (left posterior and right posterior) to better summarise the overall topography. **(c)** Distribution of all feature importances in the feature set.

## 4. Discussion

In this study, we demonstrate that psychotic-like experiences in the healthy population can be predicted from EEG data using machine learning. Specifically, we show that the brain activity evoked in response to repeating sounds within a stable environment is predictive of psychotic-like experiences at the individual level.

In our spatiotemporal models, we found that the brain responses to highly repetitive or ‘standard’ sounds, were more predictive of PQ scores than the deviant and differential MMN responses. Within the corresponding feature importance maps, voxels contributing most to lower predicted scores lie primarily within the P50 and P200 time windows of higher predicted scores concentrated within the N100 window, in keeping with known alterations in schizophrenia (Rosburg et al., 2008) and genetically high-risk groups (Larsen et al., 2018). Failure to suppress the P50 and N100 have consistently been reported in the schizophrenia literature, not only in chronic schizophrenia but also in at-risk groups (Brockhaus-Dumke et al., 2008) and relatives of patients (Turetsky et al., 2008), linking these deficits as potential endophenotypes for schizophrenia (Earls et al., 2016). Such findings are in keeping with the predictive coding theory which considers these early effects as a failure to encode the patterns associated with the standard sound (Adams et al., 2013). As a result, all stimuli, both standard and deviant sounds, are rendered surprising. Consequently, aberrant predictive models may be formed and what should be predictable sensory stimuli may appear unusually salient, leading to the formation of unusual percepts and beliefs. The P200 has also been reported as reduced in schizophrenia patients (Ford et al., 2016), particularly in response to standard tones (Ferreira-Santos et al., 2012). One possible interpretation is that surprise responses to oddball sounds irrespective of volatility are similar for all individuals, regardless of their psychotic-like experiences. The same could be said about surprise responses to standard sounds under volatile conditions since what the next sound will be is harder to predict. While a standard sound should be predicted in this scenario, the precision of such a prediction is much smaller than in less volatile environments. In stable environments, standard sounds should be more precisely predicted, provided the individual is able to form a good statistical representation of its environment. However, in schizophrenia, such model representations are thought to be less precise (Sterzer et al., 2018). As a consequence, events that should be predicted, and hence downplayed, are overly surprising by virtue of being misattributed with more salience than appropriate. Assuming that varying degrees of salience misattribution also align on a continuum and are expressed in the brain responses to precisely predictable (or stable) standards, then it is possible that such responses are discriminable of psychotic-like experiences in healthy individuals, as observed here.

As mentioned above, the analyses presented here have been repeated for a second time with increased sample size. Initially, our feature selection approach had suggested that DCM parameters, in particular, the right IFG to right STG connection were predictive of psychotic-like experience, however, this was ultimately found to be a false positive result. For the spatiotemporal approach, we had also identified seemingly significant results for all feature sets within the stable condition. Of these, only the stable standard model remained significant following the second round of testing and increased sample size, albeit with reduced performance.

Technical challenges associated with this type of modelling stem from the fact that in typical neuroimaging applications, the number of available features in high-dimensional data informing model predictions vastly outweighs the number of samples (Bellman, 1961). As such, fitting a continuous function to a high-dimensional, yet sparsely populated space which also generalises to new samples becomes difficult. In order to control for any feature selection bias and overfitting model parameters (Arbabshirani et al., 2017), we applied two different methodologies. Firstly, we extracted sets of features from multiple domains *a priori*, then using sequential feature selection, we were able to vastly reduce the number of features and pinpoint those which contributed most to the model. We then utilised an existing and widely used machine learning method in neuroimaging known as kernel learning (Schölkopf and Smola, 2000), which instead reduces the risk of overfitting by computing pairwise similarities between images and further constrains the size of feature weights through regularisation. With this approach, we obtained a spatiotemporal representation of the predictive function, indicating which brain regions and time points were assigned the largest weighted contributions in predicting the outcome variable (Fernandes Jr et al., 2017; Portugal et al., 2016), however direct interpretation of these weight maps is not as straightforward (Haufe et al., 2014; Schrouff et al., 2018).

Whilst the spatiotemporal stable standard model was statistically significant, the corresponding ERP component used in the sequential feature routine was unable to predict the true scores. One explanation for the discrepancy in performance between these two approaches is that the ERP components only represent a single datapoint in space and time, extracted from a channel and time window defined *a priori* as where the prediction error signal was thought to be most robust. Conversely, the spatiotemporal model uses the whole data, hence more powerful, and does not make any such assumptions about when or where differences are likely to occur.

All cross-validation schema commonly used to assess model generalisability have an inherent bias and variance in their error estimation. Throughout our analysis, we have been conscious of limiting potential biases in our sampling and thus estimated model performance using a 10-fold cross-validation scheme with 10 repetitions. Varoquaux recently demonstrated that commonly used schema in neuroimaging studies may report an inflated level of performance (Little et al., 2017; Varoquaux et al., 2017), namely leave-one-out cross-validation (Veronese et al., 2013). Although it is commonplace in neuroimaging to compute a single cross-validation split, the broader field of machine learning typically perform repeated cross-validation schema with varied sampling, averaging the results across replicates (Witten and Frank, 2005), which is demonstrably more robust to random variations in the training and test partitioning (Rodriguez et al., 2010). By employing these methods here, our reported performance metrics are thus more conservative estimates of model generalisability.

In this study we have used the total PQ score as the target variable in its raw form as defined in the original psychometric instrument (Loewy et al., 2005). By using one model to predict the compound score, this could be considered an early integration of the target variables given it is the sum of four constituent subscales. Another approach would be to train an ensemble of models, each on an individual subscale and performing a late integration of the model outputs to obtain the total score (Taylor et al., 2020). Whilst these approaches would be more specific and assist in clinical interpretation, this may compromise signal detection and model sensitivity.

There are a number of key limitations in this study. The PQ and our screening protocols are based on self-report, therefore we are reliant on participants recollection of past experiences and transparency in sharing information (for example, their use of illicit substances). In both cases, this introduces noise into the target variables. To a lesser extent, we are also reliant on accurate self-report of tobacco and caffeine intake, which can introduce noise to the EEG signal. Another limitation is that the sample presented here only comprises high functioning individuals who do not require clinical assistance. Given that differences in auditory oddball processing have also been observed in populations without psychosis but elevated psychopathology (Kim et al., 2020), further research is required to confirm whether these effects can be used to predict the generalist notion of psychotic experiences in higher severity, clinical samples.

## 5. Conclusion

This proof-of-concept study demonstrates that spatiotemporal EEG profiles are predictive of psychotic-like experiences on a continuum within the healthy general population. The machine learning regression methods employed here go a step beyond correlation analysis, in that they can make such predictions on an individual, rather than group basis. We envisage that with further development and integration with other instruments as part of a holistic strategy (McMillan, 2016), these neuroimaging applications may have a future role to play in identifying the underlying mechanisms and ultimately augmenting the role of the clinical practitioner.

## Acknowledgements

We would like to thank Clare Harris and Moritz Bammel for contributions toward data collection and Roshini Randeniya for assistance with dynamic causal modelling analysis.

