## Supplemental Material for "Predicting subclinical psychotic-like experiences on a continuum using machine learning"

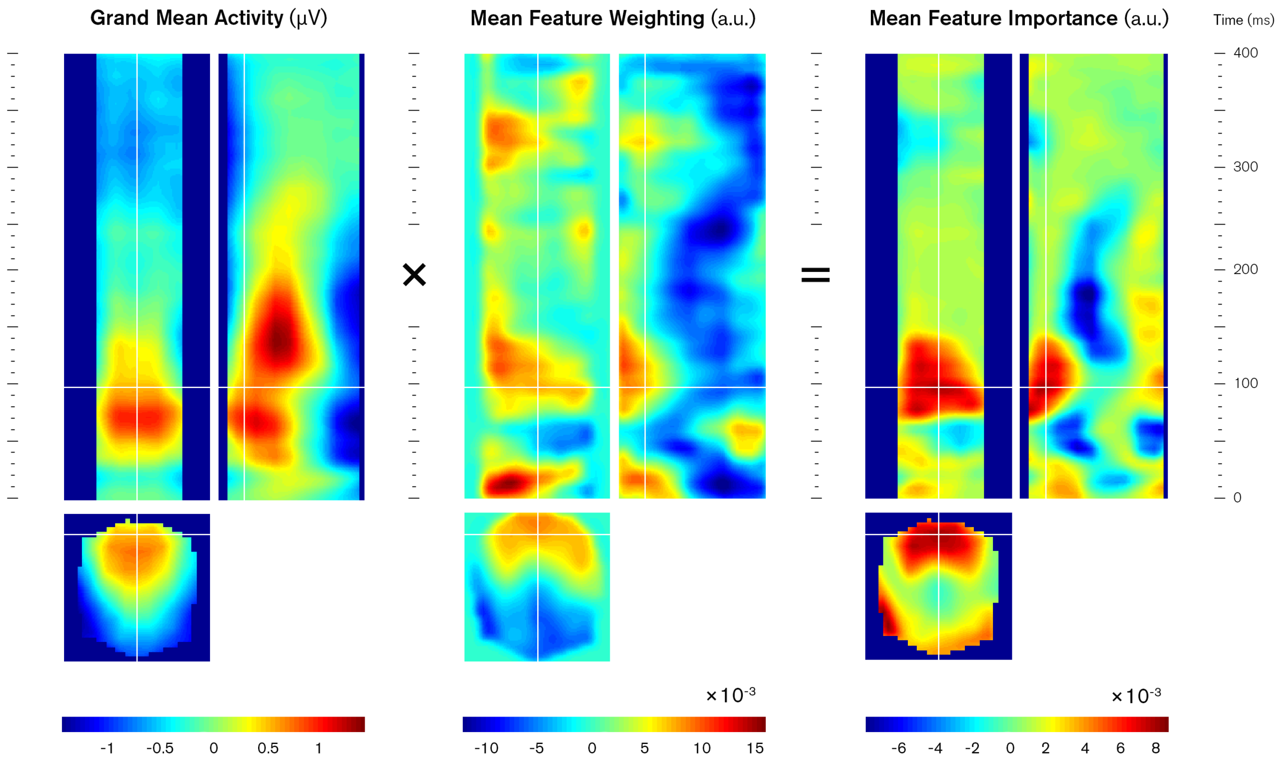


FIGURE S1 — Exemplar orthogonal views of spatiotemporal feature importance calculation (right) as the element-wise multiplication of the grand mean activity (left) and feature weights (centre). White lines indicate the intersection of spatiotemporal planes through a point of high importance in predicting high PQ scores (red), occurring frontally at approximately 100ms post-stimulus. Scalp maps orientations are annotated left (L), right (R), anterior (A) and posterior (P).

TABLE S1 — Summary of initial model performance. *p*-values were computed via 1000 permutations with Bonferroni correction for multiple comparisons.

| Method | Features | R | R *p*-value (uncorrected) | R *p*-value (corrected) | MSE | MSE *p*-value (uncorrected) | MSE *p*-value (corrected) |
|---|---|---|---|---|---|---|---|
| Feature Extraction and Selection | Behavioural | -0.02 | 0.528 | 1.000 | 1.514 | 0.528 | 1.000 |
| | ERP Components | 0.15 | 0.210 | 0.840 | 1.488 | 0.210 | 0.840 |
| | DCM Parameters | 0.43 | 0.012 | 0.048 | 1.267 | 0.012 | 0.048 |
| | Combined | 0.10 | 0.330 | 1.000 | 1.806 | 0.322 | 1.000 |
| Spatiotemporal Images | Stable Standard | 0.58 | 0.001 | 0.008 | 0.963 | 0.001 | 0.008 |
| | Stable Deviant | 0.33 | 0.002 | 0.016 | 1.301 | 0.005 | 0.040 |
| | Stable MMN | 0.49 | 0.001 | 0.008 | 1.095 | 0.001 | 0.008 |
| | Stable Combined | 0.46 | 0.001 | 0.008 | 1.128 | 0.002 | 0.016 |
| | Volatile Standard | 0.33 | 0.007 | 0.056 | 1.298 | 0.004 | 0.032 |
| | Volatile Deviant | 0.22 | 0.017 | 0.136 | 1.410 | 0.104 | 0.832 |
| | Volatile MMN | 0.01 | 0.259 | 1.000 | 1.622 | 0.987 | 1.000 |
| | Volatile Combined | 0.27 | 0.010 | 0.080 | 1.351 | 0.013 | 0.104 |

—

FIGURE S2 *(over)* — Feature selection matrices and weight vectors for each feature set, indicating the frequency at which each feature was selected and the contributions to initial model predictions upon selection. Matrices were computed as the average across all $K$ folds and $R$ repetitions. The mean selection rate and weightings are the average across all feature counts, i.e. 1 through $M$. The optimal feature count, $m$, for each model is highlighted with red dotted line. Feature selection rates are indicated using warm colormap and feature weightings are shown in greyscale.

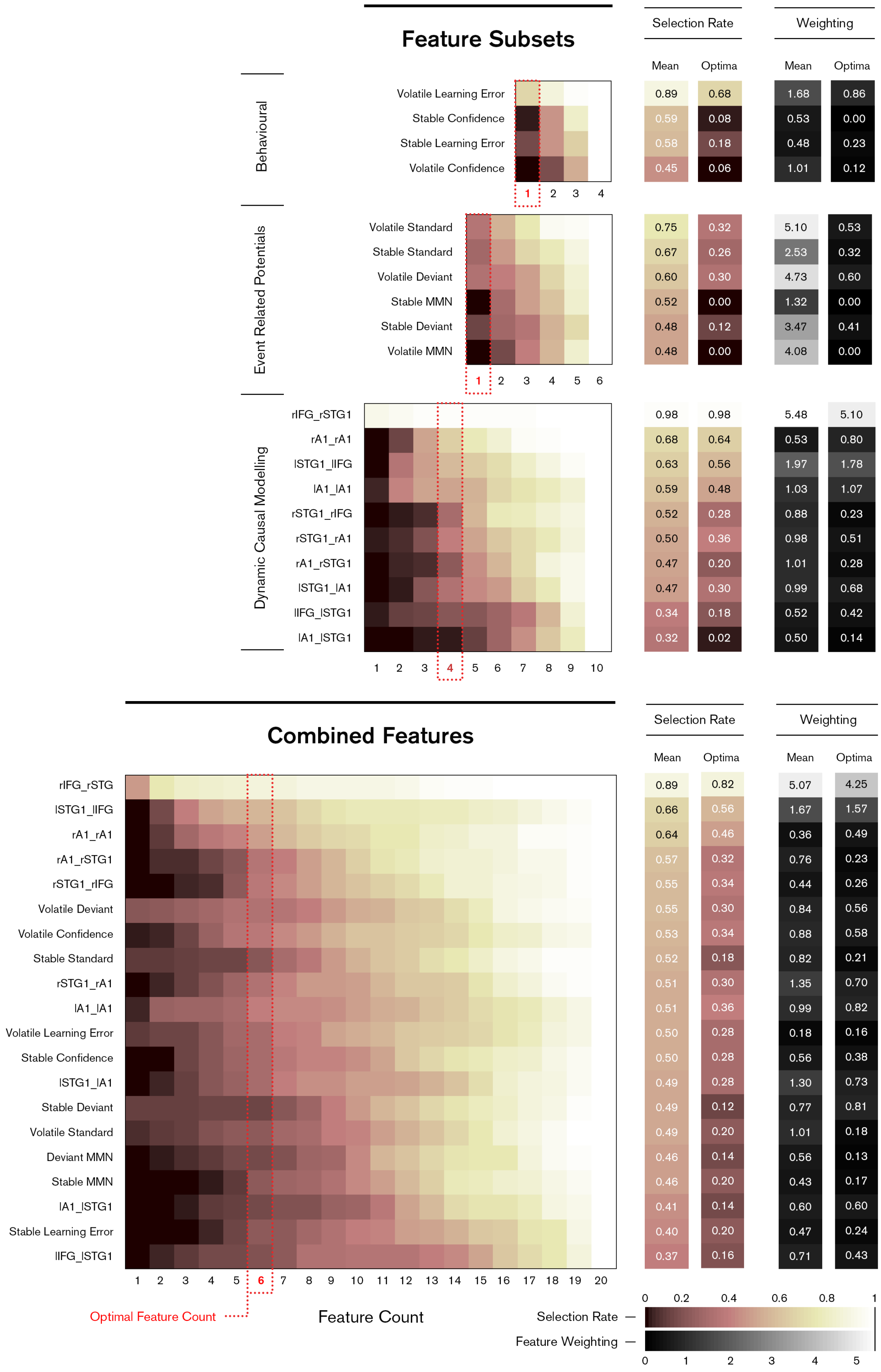

Feature Subsets
Selection Rate
Weighting
Mean
Optima
Mean
Optima
Behavioural
Volatile Learning Error 0.89 0.68 1.68 0.86
Stable Confidence 0.59 0.08 0.53 0.00
Stable Learning Error 0.58 0.18 0.48 0.23
Volatile Confidence 0.45 0.06 1.01 0.12
1 2 3 4
Event Related Potentials
Volatile Standard 0.75 0.32 5.10 0.53
Stable Standard 0.67 0.26 2.53 0.32
Volatile Deviant 0.60 0.30 4.73 0.60
Stable MMN 0.52 0.00 1.32 0.00
Stable Deviant 0.48 0.12 3.47 0.41
Volatile MMN 0.48 0.00 4.08 0.00
1 2 3 4 5 6
Dynamic Causal Modelling
rIFG_rSTG1 0.98 0.98 5.48 5.10
rA1_rA1 0.68 0.64 0.53 0.80
lSTG1_lIFG 0.63 0.56 1.97 1.78
lA1_lA1 0.59 0.48 1.03 1.07
rSTG1_rIFG 0.52 0.28 0.88 0.23
rSTG1_rA1 0.50 0.36 0.98 0.51
rA1_rSTG1 0.47 0.20 1.01 0.28
lSTG1_lA1 0.47 0.30 0.99 0.68
lIFG_lSTG1 0.34 0.18 0.52 0.42
lA1_lSTG1 0.32 0.02 0.50 0.14
1 2 3 4 5 6 7 8 9 10
Combined Features
Selection Rate
Weighting
Mean
Optima
Mean
Optima
rIFG_rSTG 0.89 0.82 5.07 4.25
lSTG1_lIFG 0.66 0.56 1.67 1.57
rA1_rA1 0.64 0.46 0.36 0.49
rA1_rSTG1 0.57 0.32 0.76 0.23
rSTG1_rIFG 0.55 0.34 0.44 0.26
Volatile Deviant 0.55 0.30 0.84 0.56
Volatile Confidence 0.53 0.34 0.88 0.58
Stable Standard 0.52 0.18 0.82 0.21
rSTG1_rA1 0.51 0.30 1.35 0.70
lA1_lA1 0.51 0.36 0.99 0.82
Volatile Learning Error 0.50 0.28 0.18 0.16
Stable Confidence 0.50 0.28 0.56 0.38
lSTG1_lA1 0.49 0.28 1.30 0.73
Stable Deviant 0.49 0.12 0.77 0.81
Volatile Standard 0.49 0.20 1.01 0.18
Deviant MMN 0.46 0.14 0.56 0.13
Stable MMN 0.46 0.20 0.43 0.17
lA1_lSTG1 0.41 0.14 0.60 0.60
Stable Learning Error 0.40 0.20 0.47 0.24
lIFG_lSTG1 0.37 0.16 0.71 0.43
1 2 3 4 5 6 7 8 9 10 11 12 13 14 15 16 17 18 19 20
Optimal Feature Count
Feature Count
Selection Rate
Feature Weighting
0 0.2 0.4 0.6 0.8 1
0 1 2 3 4 5

FIGURE S3 — Plot of initial model predictions and true observed PQ scores. (A) DCM connectivity feature set, using Huber regression and forward selection, optimised to four features; (B) Spatiotemporal stable standard ERP feature set, using kernel ridge regression and regularisation. Predictions from each repetition are shown as grey dots and overall predictions as black crosses. All scores are normalised via log transform.

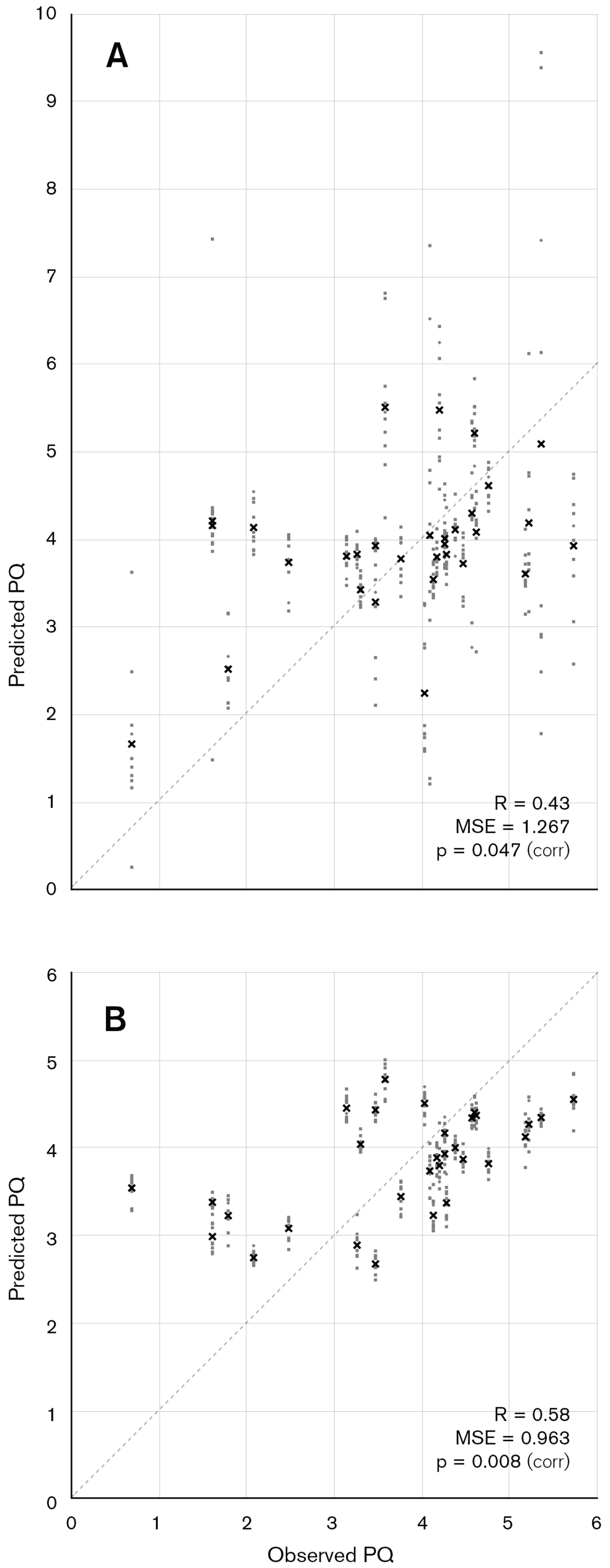

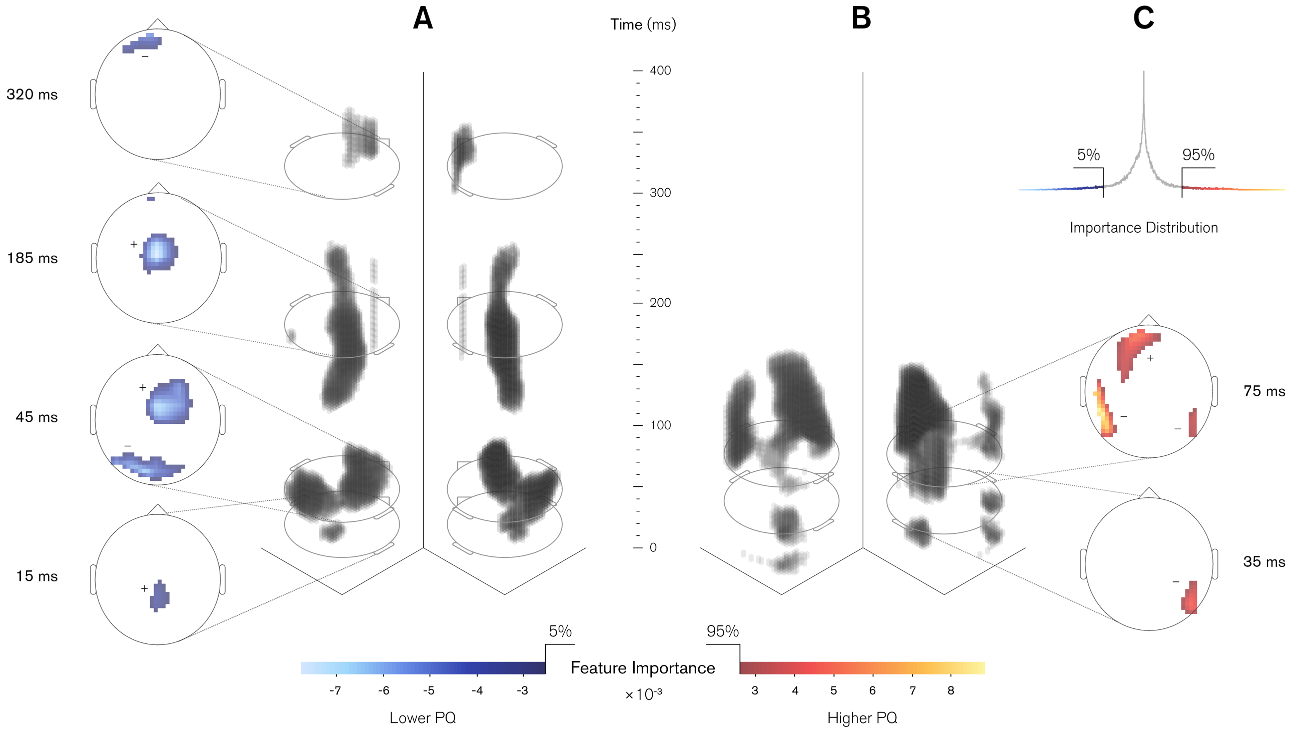


FIGURE S4 — Spatiotemporal features of greatest importance to schizotypy predictions in initial stable standard response model. (A-B) 3D volumes represent clusters of greatest feature importance for low (cool) and high (warm) predictions of PQ, thresholded at top 5% of negative and positive importances respectively, with spatial dimensions on *x*-*y* plane and time domain along vertical *z*-axis, displayed from dual angles. 2D scalp maps refer to peaks within main clusters, with ± labels indicating polarity of the original ERP signal; (C) Distribution of all feature importances in the feature set.
